# Mechanistic Dissection of Spatial Organization in NF-κB Signaling Pathways by Hybrid Simulations

**DOI:** 10.1101/2020.11.11.378331

**Authors:** Yinghao Wu, Kalyani Dhusia, Zhaoqian Su

**Affiliations:** Department of Systems and Computational Biology, Albert Einstein College of Medicine, 1300 Morris Park Avenue, Bronx, NY, 10461

## Abstract

The nuclear factor kappa-light-chain-enhancer of activated B cells (NF-κB) is one of the most important transcription factors involved in the regulation of inflammatory signaling pathways. Inappropriate activation of these pathways has been linked to autoimmunity and cancers. Emerging experimental evidences have been showing the existence of elaborate spatial organizations for various molecular components in the pathways. One example is the scaffold protein tumor necrosis factor receptor associated factor (TRAF). While most TRAF proteins form trimeric quaternary structure through their coiled-coil regions, the N-terminal region of some members in the family can further be dimerized. This dimerization of TRAF trimers can drive them into higher-order clusters as a response to receptor stimulation, which functions as a spatial platform to mediate the downstream poly-ubiquitination. However, the molecular mechanism underlying the TRAF protein clustering and its functional impacts are not well-understood. In this article, we developed a hybrid simulation method to tackle this problem. The assembly of TRAF-based signaling platform at the membrane-proximal region is modeled with spatial resolution, while the dynamics of downstream signaling network, including the negative feedbacks through various signaling inhibitors, is simulated as stochastic chemical reactions. These two algorithms are further synchronized under a multiscale simulation framework. Using this computational model, we illustrated that the formation of TRAF signaling platform can trigger an oscillatory NF-κB response. We further demonstrated that the temporal patterns of downstream signal oscillations are closely regulated by the spatial factors of TRAF clustering, such as the geometry and energy of dimerization between TRAF trimers. In general, our study sheds light on the basic mechanism of NF-κB signaling pathway and highlights the functional importance of spatial regulation within the pathway. The simulation framework also showcases its potential of application to other signaling pathways in cells.

## Introduction

The innate immune system constitutes the first line of host defense during infections by invading pathogens [1]. This defensive response, called inflammation, is a complicated process orchestrated by many different cellular components [2, 3]. At the onset of inflammation, cytokines are released from immune cells such as macrophage after they capture infected cells [2]. The nuclear factor kappa-light-chain-enhancer of activated B cells (NF-κB) is a critical transcription factor regulates the expressions of cytokines that stimulate inflammatory responses [4, 5]. Under normal condition, NF-κB is retained in the cytoplasm with an inhibitory factor, IκB [6, 7]. The NF-κB activation is started from the binding of membrane receptors, mainly in tumor necrosis factor receptor (TNFR) superfamily to their extracellular ligands [8]. The ligand binding of receptors triggers the recruitment of adaptor proteins, such as TNF receptor associated factor (TRAF), to their cytoplasmic domains [9]. While the C-terminal domain of TRAF maintains interactions with upstream receptors, the N-terminal regions function as platform to mediate the process of poly-ubiquitination [10]. The ubiquitination leads to the degradation of IκB, so that NF-κB can be released and enter cell nucleus to initiate gene expression [11, 12]. It has been found that spatial organizations of proteins are highly involved in signaling pathways [13, 14]. Such phenomena have also been observed in NF-κB signaling pathway [15]. For instance, *in vivo* experiments have shown that TRAF proteins can aggregate to higher-order spatial patterns as a response to receptor stimulation [16, 17]. The structural evidences further indicate that the N-terminal region in some TRAF proteins, such as TRAF6 and TRAF2, is dimeric [16, 18, 19]. The capability of high-order pattern formation can be abolished by a mutant that disables this dimerization. As a result, it was proposed that the dimerization can lead to a two-dimensional clustering of trimeric TRAF protein complexes. However, it is not fully understood how this spatial organization regulates the dynamics in NF-κB signaling pathway.

Due to the functional importance of NF-κB in immunity, the molecular mechanisms of its signaling pathway are under intensive investigation [20]. Comparing with other traditional wet-lab experiments, computational modeling is more convenient to explore the complexity of a biological system on a mechanistic level. As a result, a large variety of models have been developed on different scales of the pathway. The original model focused on the role of IκB in regulating the temporal dynamics of NF-κB by using a set of ordinary differential equations (ODE) [21, 22]. More recently, stochastic simulations and agent-based modeling (ABM) approaches were applied to generate new hypotheses on the behavior of various molecular components in the NF-κB pathway [23-26]. However, information about spatial organization of these components, such as the oligomerization of TRAF proteins, has not been incorporated into these models. On the other hand, methods in another class of simulation technique, including MCELL [27], Smoldyn [28, 29], and more recently SpringSaLaD [30, 31], use lattice-based or particle-based models to implement the diffusions of biomolecules within a more realistic cellular environment. They can be used to simulate the formation of spatial patterns on the subcellular level, such as protein complex assembly or phase separation [32-40]. Unfortunately, these methods are intractable to be applied to study the dynamics of an entire signaling pathway because of their high demands for computational resources. Overall, it is highly challenging to develop a simulation method which is able to capture a signaling event with both spatial resolution and its functional impacts on the whole signaling networks.

In order to overcome this challenge, a hybrid simulation approach is presented to describe the dynamics in the NF-κB signaling pathway. The method contains two coupled systems. The assembly of TRAF-based signaling platform at the membrane-proximal region is modeled by a rigid-body-based diffusion-reaction algorithm, while the downstream signaling events, including the recruitment of IκB kinase (IKK), phosphorylation of IκB and activation of NF-κB are simulated as stochastic chemical reactions. These two levels of simulations are integrated together, so that how spatial variations in the signaling platform affect the rest of the pathway can be quantitatively analyzed. The signaling network is further regulated by negative feedbacks through different inhibitors such as IkB and A20, which leads to an oscillatory NF-κB response. We demonstrated that the temporal patterns of downstream signal oscillations are closely correlated with the spatial organization of upstream platform assembly. Our simulation results also suggested that the cellular dynamics of NF-κB signaling pathway can be fine-tuned by a single pair of molecular interaction between TRAF proteins. Altogether, this study sheds light on the basic mechanism of NF-κB signaling pathway and highlights the functional importance of spatial regulation. The hybrid simulation framework also showcases its potential of application to other systems of cell signaling pathways.

## Results and Discussions

### General description of the outputs from simulations of the signaling network

Most members in TRAF family contain a RING domain at their N-terminus, a TRAF-C domain with seven to eight anti-parallel β-strand folds at their C-terminus, and a coiled-coil region in the middle [41]. While the C-terminal domain of TRAF maintains interactions with upstream receptors, the N-terminal regions function as platform to mediate the downstream poly-ubiquitination, as well as the *cis*-interactions between themselves [18]. Moreover, most members in TRAF family form trimeric quaternary structure through their coiled-coil regions [42]. We are mainly focusing on the spatial clustering of TRAF trimers and its impacts on regulating the downstream signaling dynamics. The formation of upstream ligand-receptor complex is beyond the scope of this study. As a result, we assume that the ligand-receptor complexes have already preformed on cell surface before our simulations and the trimeric scaffold proteins have already bound to the cytoplasmic domains of receptors (**Figure 1a**). Based on this assumption, the TRAF trimer and the ligand-receptor complex are modeled as one single rigid body attached to the cell membrane. Each rigid body contains three binding sites to maintain the *cis*-interaction between TRAF trimers. The movements of each TRAF trimer are confined within the two-dimensional membrane proximal area. The multiple *cis*-interactions involved in each TRAF trimer and its two-dimensional movements can further result in the higher-order clustering, which is simulated by a diffusion-reaction algorithm. Detailed model representation of TRAF trimers and the follow-up algorithm used to simulate their clustering are described in the **Methods**.

**Figure 1:**
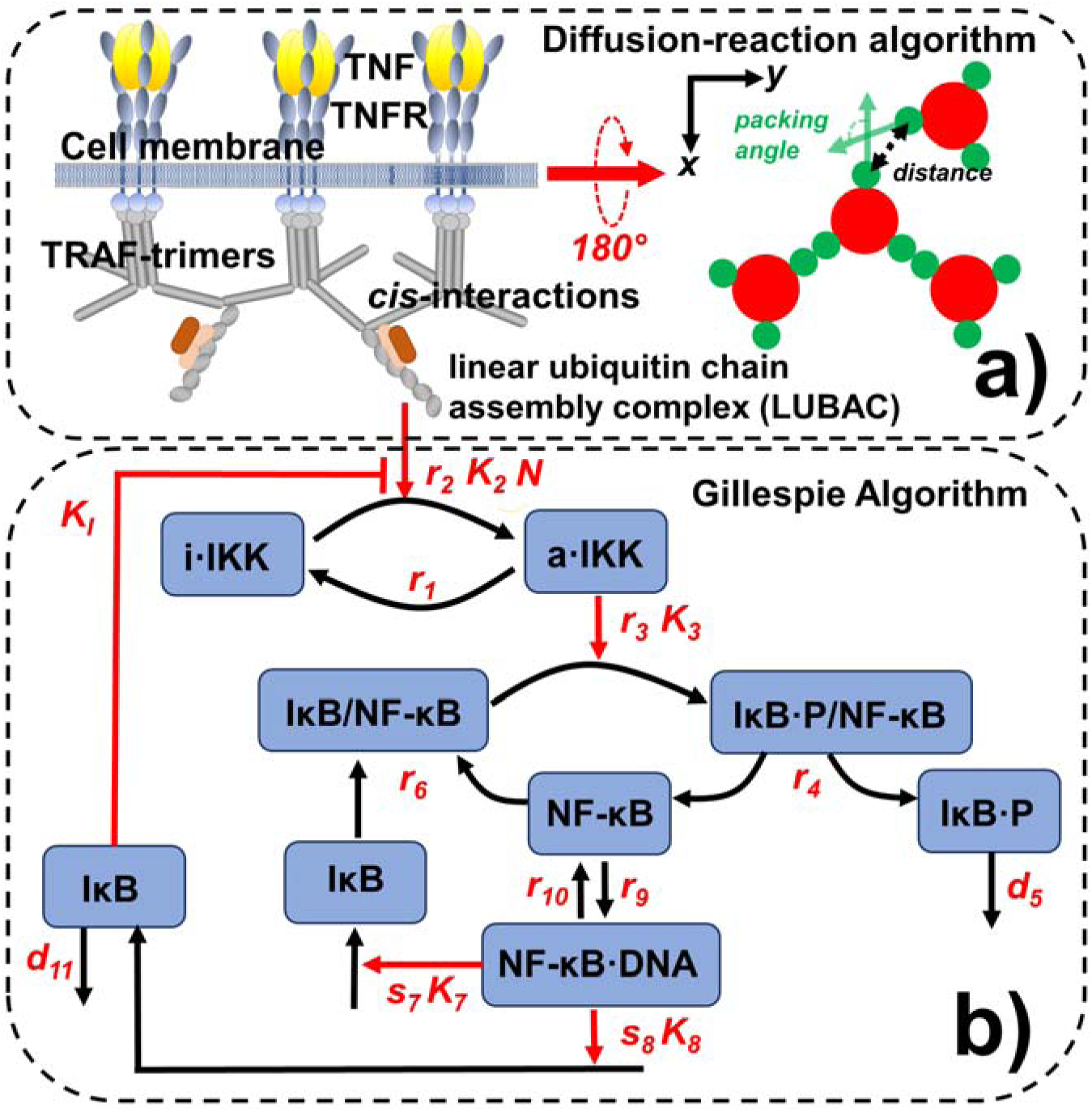
The dynamics of the NF-κB signaling pathway is simulated by a computational method consisting of two coupled systems. The assembly of TRAF-based signaling platform at the membrane-proximal region is modeled by a rigid-body-based diffusion-reaction algorithm, as shown in **(a)**. We assume that each TRAF trimer has bound to its upstream ligand-receptor complex and therefore is modeled as a single rigid body which movements are confined within the two-dimensional membrane proximal area. The formation of a *cis*-interaction between two TRAF trimers thus facilitates the assembly of linear ubiquitin chain assembly complex (LUBAC). The LUBAC further provide the scaffold to enable the activation of downstream signaling pathway, which diagram is sketched in **(b)**. The dynamics of the signaling network is simulated by Gillespie algorithm. Finally, the diffusion-reaction and Gillespie algorithms are synchronized under a multiscale simulation framework, as described in the **Method**.

In addition to form *cis*-interactions, the N-terminal RING domain of TRAF protein also functions as an E3 ubiquitin ligase. It recruits the E2 ubiquitin-conjugating enzyme such as Ubc13 so that the poly-ubiquitin chains can be formed [19]. The formation of a *cis*-interaction between two TRAF trimers thus facilitates the assembly of linear ubiquitin chain assembly complex (LUBAC) [43]. The LUBAC further provide the scaffold to enable the activation of the kinases IKK. Upon activation, IKK can induce the phosphorylation of IκB, which forms a complex with the transcription factor NF-κB as an inhibitor [11]. The complex will be dissociated after the phosphorylation of IκB, so that the transcription factor can be freely released. The phosphorylated inhibitor later undergoes ubiquitin-dependent proteolysis, while the released transcription factor enters cell nucleus and bind to the promoter and turn on its target genes. The genetic regulation of two proteins is specific modeled in the network. The first is the inhibitor IκB itself. The newly synthesized IκB enters cytoplasm and associates with free NF-κB to block its function as a transcription regulator [44]. The second protein which gene is also turned on by NF-κB is A_20_. Upon synthesis, the protein also functions as an inhibitor to diminish the activation process of IKK [45]. The diagram of the network is shown in **Figure 1b**, while its mathematical description is presented in the **Method**.

Based on the mathematical representation of all reactions in the signaling network, the dynamics of the system is simulated stochastically by Gillespie algorithm. The signaling network has further been coupled with the spatial clustering of TRAF trimers under a hybrid simulation framework. The specific procedure of the hybrid simulation which combined Gillespie algorithm with the spatial model of TRAF clustering is delineated in the **Method**. The results from this hybrid simulation are summarized in **Figure 2**. Specifically, 200 TRAF trimers were randomly placed in the two-dimensional membrane proximal region as an initial configuration (**Figure 2a**). The length of each side in this square region is 1000nm, along both X and Y directions, which gives a total area of 1µm^2^. Some representative snapshots of the clustering process are also plotted along the simulation trajectory. We found that TRAF trimers start to aggregate into small oligomers (**Figure 2b**). These oligomers are organized into hexagonal lattice based on the spatial symmetry of three binding sites in each trimer. The formation of these oligomer is a very dynamic process. TRAF trimers constantly left one oligomer and joined another one. As a result, small oligomers either disappeared or merged into neighboring larger oligomers, leading into a configuration in which the number of clusters became smaller and their sizes continued growing, as shown in **Figure 2c**. Ultimately, most trimers aggregated together into a final larger cluster and the system reached equilibrium, as shown in **Figure 2d**.

**Figure 2:**
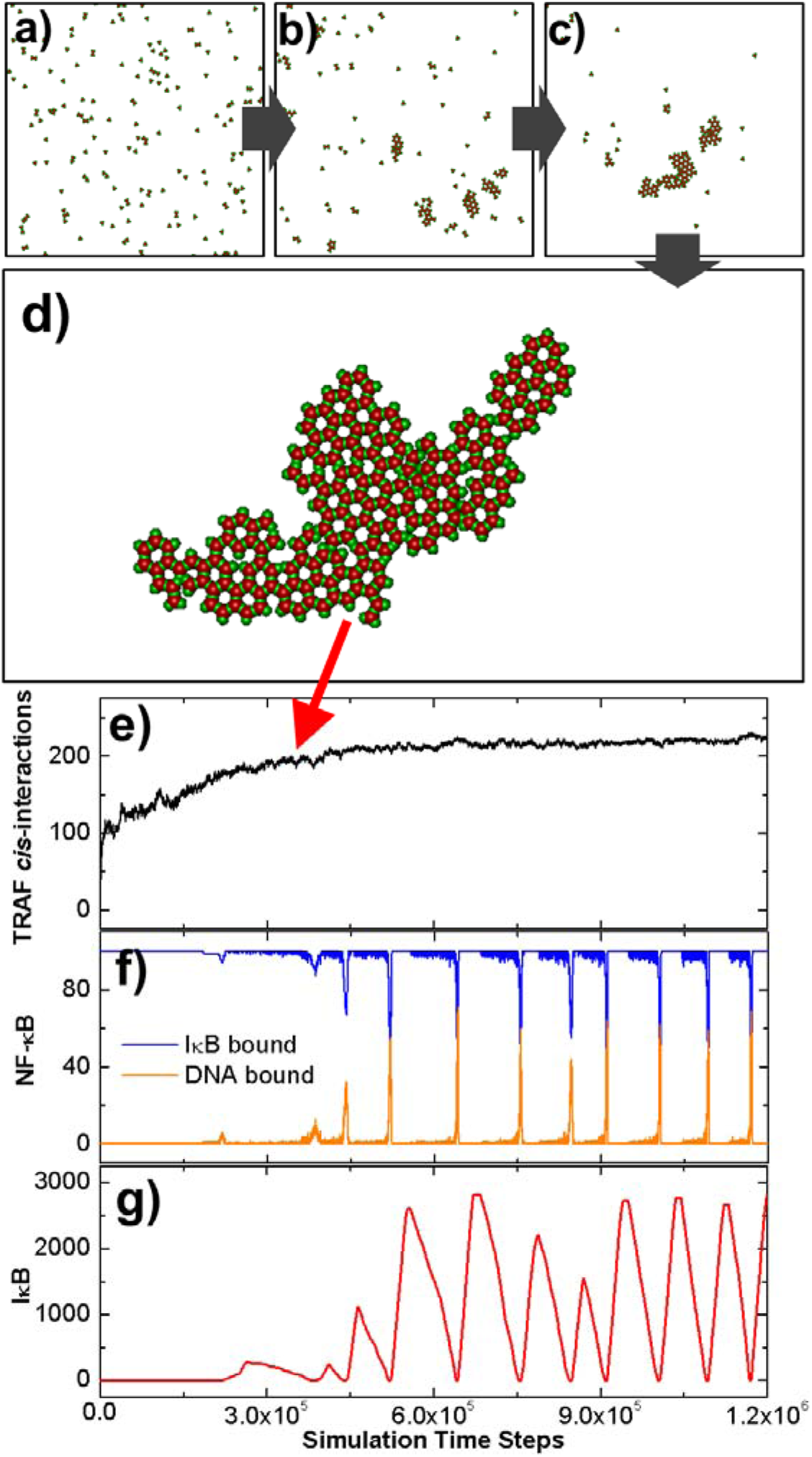
The initial configuration of the diffusion-reaction simulation is shown in **(a)**. Some representative snapshots along the simulation trajectory were selected in **(b)** and **(c)**. The configuration that the system reached equilibrium is further shown in **(d)**. In **(e)**, we plotted the number of *cis*-interactions between TRAF trimers observed in the system along with the simulation time steps. The kinetic profiles of molecular components in the downstream signaling network were further plotted as functions of simulation time steps. These profiles are: the numbers of NF-κB molecules which bind to IκB and DNA as shown by the blue and orange curves in **(f)**; and the total number of free IκB in the system as shown by the red curve in **(g)**.

The number of *cis*-interactions formed between TRAF trimers was further plotted in **Figure 2e** as a function of simulation time steps. The figure shows that the number of *cis*-interactions increased very fast at the beginning of the simulation, indicating an initial seeding process. The increase of *cis*-interactions became slower as the oligomers in the system grew, and finally reached saturation after 4.5×10^5^ simulation time steps, corresponding to the formation of the final cluster. The kinetic profiles of molecules in the downstream signaling network were further plotted in the following figures. Specifically, the numbers of NF-κB molecules which bind to IκB and DNA are shown in blue and orange curves in **Figure 2**.**f**, respectively. A periodic boost of DNA-bound NF-κB is displayed in the figure, corresponding to the timing of dissociation between NF-κB and IκB. Following the cycle of NF-κB, the total number of free IκB in the system also oscillated along the simulation, as illustrated in **Figure 2**.**g**. This oscillatory behavior is the result of the negative feedback regulation in the network, which is implemented by IκB and A_20_. Additionally, we found that the oscillation was initiated after the number of total *cis*-interactions between TRAF trimers almost stop changing, indicating that the formation of large clusters is critical to activate the downstream signaling pathway. In summer, our simulation results showcased the oscillatory dynamics in the NF-κB signaling pathway and its correlation with the spatial organization of molecular components in the pathway.

### The comparison of signaling dynamics in systems with and without TRAF clustering

In order to further elucidate the functional importance of TRAF clustering in regulation of downstream signaling pathway, we carried out two separate simulations with different scenarios. The TRAF trimers in the first scenario, as we descried before, contain three binding sites. As a result, each trimer can simultaneously form *cis*-interactions with three structural neighbors, leading into the formation of a highly ordered spatial pattern. In contrast, a control system was designed in the second scenario. In this control system, we disabled the possibility of high-order clustering among TRAF proteins. This was achieved by assigning only one binding site to each TRAF protein instead of three, so that only *cis*-dimers can be formed between two proteins. As the initial configuration of the first scenario, 200 rigid bodies of TRAF trimers were randomly placed on a two-dimensional square surface with an area of 1000×1000 nm^2^. On the other hand, the control simulation contains 600 rigid bodies randomly distributed on the surface with the same area to maintain the same level of possible total interactions as in the first scenario. All the other parameters in the diffusion-reaction simulation of control scenario such as diffusion and binding constants remain unchanged. The binding affinity of *cis*-interactions in both systems equals -10kT. Moreover, in the stochastic simulations of downstream signaling network, the same values of rate constants were used for the second scenario as for the first scenario.

The comparison of simulation results between these two scenarios are summarized in **Figure 3**. The total number of *cis*-interactions formed between TRAF trimers along the simulation in the first system is plotted by the red curve in **Figure 3**.**a**, while the total number of dimerized *cis*-interactions formed in the control system is plotted by the red curve. The figure shows that the number of *cis*-interactions in the system which can form clusters grew more slowly but reached a much higher level than the system which can only form dimers, although the binding affinity and total binding sites in both systems are the same. Moreover, the level of fluctuations in the first system is also lower than the second one. The final configurations at the end of two simulations are compared with each other in **Figure 3c** and **Figure 3d**, respectively. Different from a small number of large clusters oligomerized in the first system, a large number of dimers are randomly distributed in the control system.

**Figure 3:**
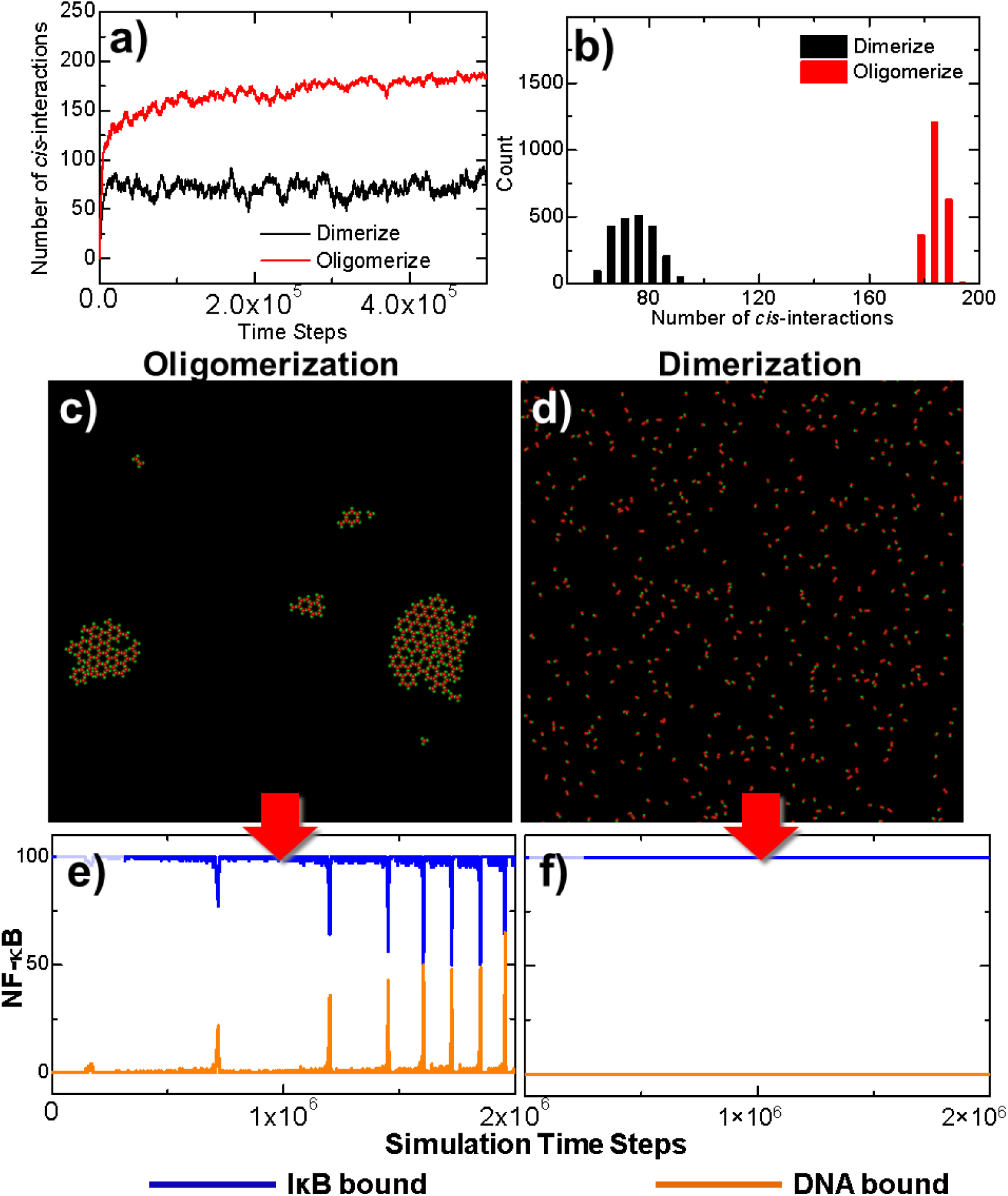
Two simulation scenarios were designed to elucidate the functional importance of TRAF clustering in regulation of downstream signaling pathway. The TRAF trimers in the first scenario contain three binding sites and can simultaneously form *cis*-interactions with three structural neighbors, while in a control system only one binding site was assigned to each TRAF protein. The numbers of *cis*-interactions formed along the simulations in these two scenarios are plotted in **(a)**. The distributions of fluctuations in the number *cis*-interactions are shown as histograms in **(b)**. The final configurations at the end of two simulation systems are compared with each other in **(c)** and **(d)**. Finally, the outputs from the downstream signaling networks of these two systems, e.g. the numbers of NF-κB molecules which bind to IκB and DNA, are also compared with each other in **(e)** and **(f)**, respectively.

In order to quantitatively estimate the statistical significance of observed differences between these two systems, a two-sample student’s t-test was carried out to the data collected from these two trajectories, which distributions are plotted as histograms in **Figure 3**.**b**. In detail, the average number of *cis*-interactions in the distribution of dimerized system is 75.51 and its standard deviation equals 7.16. Relatively, the average number of *cis*-interactions in the distribution of oligomerized system is 182.53 and its standard deviation equals 3.04. The null hypothesis is that no difference exists between two distributions. It was tested at a 95% confidence interval. The calculated t-score equals 650.28 with the P-value lower than 0.0001. This small P-value from the t-test indicates that the null hypothesis can be rejected and the alternative hypothesis can be accepted, i.e., the spatial arrangement of TRAF oligomers lead into significant change in the level of *cis*-interactions comparing with the system in which oligomers are not allowed to form.

The outputs from the downstream signaling networks of the two comparative systems are shown in the following panels of the figure. The changes of NF-κB that bound to IκB or DNA in the first system are plotted as a function of the simulation time with blue and yellow curves in **Figure 3e**. The oscillations were observed in the system, in which the number of NF-κB/DNA complexes was boosted periodically after the clusters of TRAF trimers were stabilized, corresponding to the red curve in **Figure 3a**. The impulse of NF-κB/DNA complexes is coincident with the drop of NF-κB/IκB complexes, which is due to the phosphorylation of IκB by activated IKK. This oscillatory dynamic of components in the network indicates that the signaling pathway was turned on by the clustering of TRAF trimers. In contrast, the number of NF-κB/IκB complexes remained at its initial level throughout the simulation in the second system. As a result, no NF-κB/DNA complex was obtained before the end of the simulation, corresponding to the curves plotted in **Figure 3f**. This result suggests that the signaling pathway could not be effectively turned on due to the less numbers and higher fluctuations of *cis*-dimers formed in the control scenario.

In summary, this comparative study demonstrated the possibility that systems with the same sets of rate parameters can evolve into very different temporal dynamics, only because of the different spatial organizations of signaling molecules. More specifically, a threshold-like behavior is controlled by the geometric arrangement of cis-interaction between TRAF scaffold proteins. Multiple binding sites in TRAF trimers lead to their high-order aggregation at membrane proximal region. Comparing with the regular dimer, each trimer in a cluster is simultaneously involved in three *cis*-interactions with its neighbor. These interlocked systems are kinetically more difficult to be dissociated. Consequently, they provide stable platform to downstream signal activation. Without this trimeric quaternary structure, on the other hand, TRAF proteins can only dimerized. Comparing with the highly ordered oligomers, these dimers are easier to be dissociated, although the binding affinity of the *cis*-interaction remains unchanged. As a result, the poly-ubiquitination facilitated by dimerized TRAF proteins alone cannot pass the threshold of downstream signal activation.

### Change the binding constants in the oligomerization

We have estimated the importance of TRAF trimers’ multiple *cis*-binding sites not only in their assembly into higher-order structures, but also in the initiation of NF-κB signaling pathway. In this section, we further explore the energetic impacts of this *cis*-interaction on spatial-temporal dynamics of the system. Specifically, the binding affinity for a given pair of binding sites between two TRAF trimers was charged to different values in the diffusion-reaction simulation. Three scenarios were tested separately. A weak binding affinity (−6kT) was used in the first scenario. In comparison, a strong binding affinity (−14kT) was used in the second scenario. In addition to these two systems, a moderate value of binding affinity (−8kT) was also tested as the third scenario. All the other parameters in both TRAF clustering section and signaling network section of the simulations are the same. The kinetic profiles generated from the simulations of these three systems are plotted in **Figure 4**.

**Figure 4:**
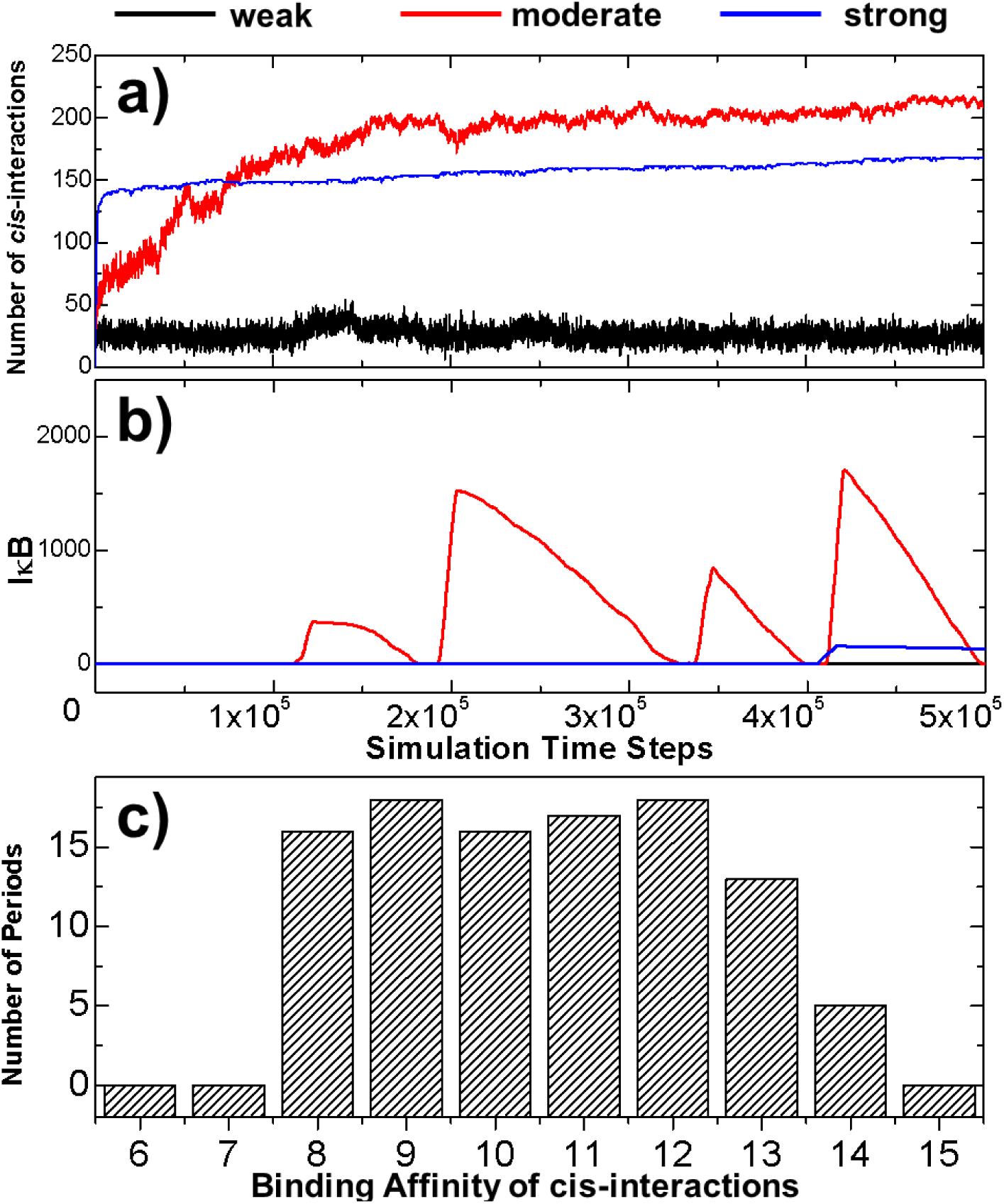
In order to explore the energetic impacts of the *cis*-interaction between TRAF trimers on spatial-temporal dynamics of the system. Three scenarios with the weak, moderate and strong binding affinities of these *cis*-interactions were tested separately. The numbers of *cis*-interactions formed in these three scenarios are plotted in **(a)** as a function of simulation time steps, while the numbers of free IκB in the system are shown in **(b)**. More systematical tests were further carried out in which we changed the binding affinities of *cis*-interaction from -6kT to -15kT with an interval of 1kT. The average numbers of oscillation periods were derived from the multiple trajectories of these system, and were plotted as histogram in **(c)** as a function of *cis*-binding affinity.

The numbers of *cis*-interactions formed in the three scenarios are plotted in **Figure 4a** as a function of simulation time steps. The profile with the weak binding affinity is shown by the black curve, which indicates a low number of *cis*-interactions. On the other hand, a much higher level of *cis*-interactions were derived in the system with strong bind affinity, as shown by the blue curve. A more interesting kinetic profile was observed in the system with moderate binding affinity, corresponding to the red curve in **Figure 4a**. The figure suggests that a small reduce of affinity from -6kT to -8kT significantly increased the level of *cis*-interactions. Surprisingly, the number of total interactions formed by the end of the simulation with moderate binding affinity is more than 200, even higher than the number found in the system with much stronger *cis*-interactions. Moreover, comparing with the system of strong affinity which reached equilibrium at the very early stage of simulation, the number of *cis*-interactions in the system of moderate binding affinity increased much slower.

The dynamics in the downstream signaling networks of three systems are further shown in **Figure 4b**. The changes of free IκB are plotted as a function of simulation time steps. An oscillation on the level of IκB was observed in the system of moderate binding affinity, as shown by the red curve in the figure. The oscillation was started when the stimulation time reached 1×10^5^ steps, consistent with the stage as the number of *cis*-interactions in the system didn’t further increase (shown by the red curve in **Figure 4a**). In contrast, no free IκB was observed in the system with weak binding affinity. This is due to the low level of *cis*-interactions formed in the system. The poly-ubiquitin chains recruited by this small number of *cis*-interaction between TRAF proteins were not enough to active IKK in the system, and prevented the IκB from being phosphorylated. As a result, the genes regulated by NF-κB cannot be turned on throughout the simulation. It is worth mentioning that no oscillation of free IκB was also obtained in the system with strong binding affinity. The IκB was produced at very late stage of the simulation after the time passed 4×10^5^ steps. However, it reached to a very low level, but then start to decay all the way through the end of the simulation, as shown by the blue curve in **Figure 4b**. The results from the simulations of these three systems therefore suggest that the oscillatory patterns of the signaling dynamics are very sensitive to the strength of *cis*-interaction between TRAF proteins.

In order to test the correlation between the oscillatory patterns of the signaling pathway and the binding affinity of the *cis*-interaction on a more statistically meaningful level, systematical tests were carried out to systems which binding affinities of *cis*-interaction were ranged from -6kT to -15kT with an interval of 1kT. Multiple trajectories were generated for each system, while a longer simulation time (2×10^6^ steps) was given for each trajectory. At the end of all simulations, we counted the number of oscillatory periods observed in each trajectory. The average number of oscillatory periods was then calculated for each system depending on the values of binding affinity. The calculated results are summarized as a histogram in **Figure 4c**. The figure confirms that oscillations were not observed under weak binding affinities, but suddenly appeared when affinities are below -7kT, suggesting that there is phase transition in the system. Further enhancement of binding affinity doesn’t change the frequency of oscillation too much until it becomes stronger than -12kT. Above this region, the oscillatory frequency will be gradually reduced if the *cis*-binding affinity increases. Finally, no more oscillation exists after the binding affinity equals -15kT. This is consistent with the signaling outputs illustrated in **Figure 4b**.

To further understand how NF-κB signals can be spatially regulated by TRAF protein clustering, the final configurations from some of the simulation systems were plotted in **Figure 5**. The final configurations from the systems which binding affinities equal -7kT and -8kT are shown in **Figure 5a** and **Figure 5b**, respectively. With only a slight change in the binding affinity, very different spatial pattern was derived in the figures. This difference can be explained by the phase transition of TRAF clustering which drives the system from a scattered distribution of trimers to their condensation into a single cluster. The network of trimers formed within the cluster stabilizes the *cis*-interactions and thus facilitate the signal activation of downstream pathway. In comparison, the final configurations from two systems selected from the other end of the binding affinity are shown in **Figure 5c** and **Figure 5d**. These figures show that systems with stronger binding affinity contain more clusters with smaller sizes. Comparing with the signal big cluster formed in the system with binding affinity of -8kT, five clusters were obtained in the system with binding affinity of -13kT and each cluster on average contains 40 trimers, as shown in **Figure 5c**. When the binding affinity reaches -15kT, as shown in **Figure 5d**, the number of clusters in the system increases to 20 and each cluster on average only contains 10 TRAF trimers. These results suggest that the strong binding affinities kinetically trap TRAF trimers into small clusters, which plays negative role in activating the oscillations in the downstream signaling network.

**Figure 5:**
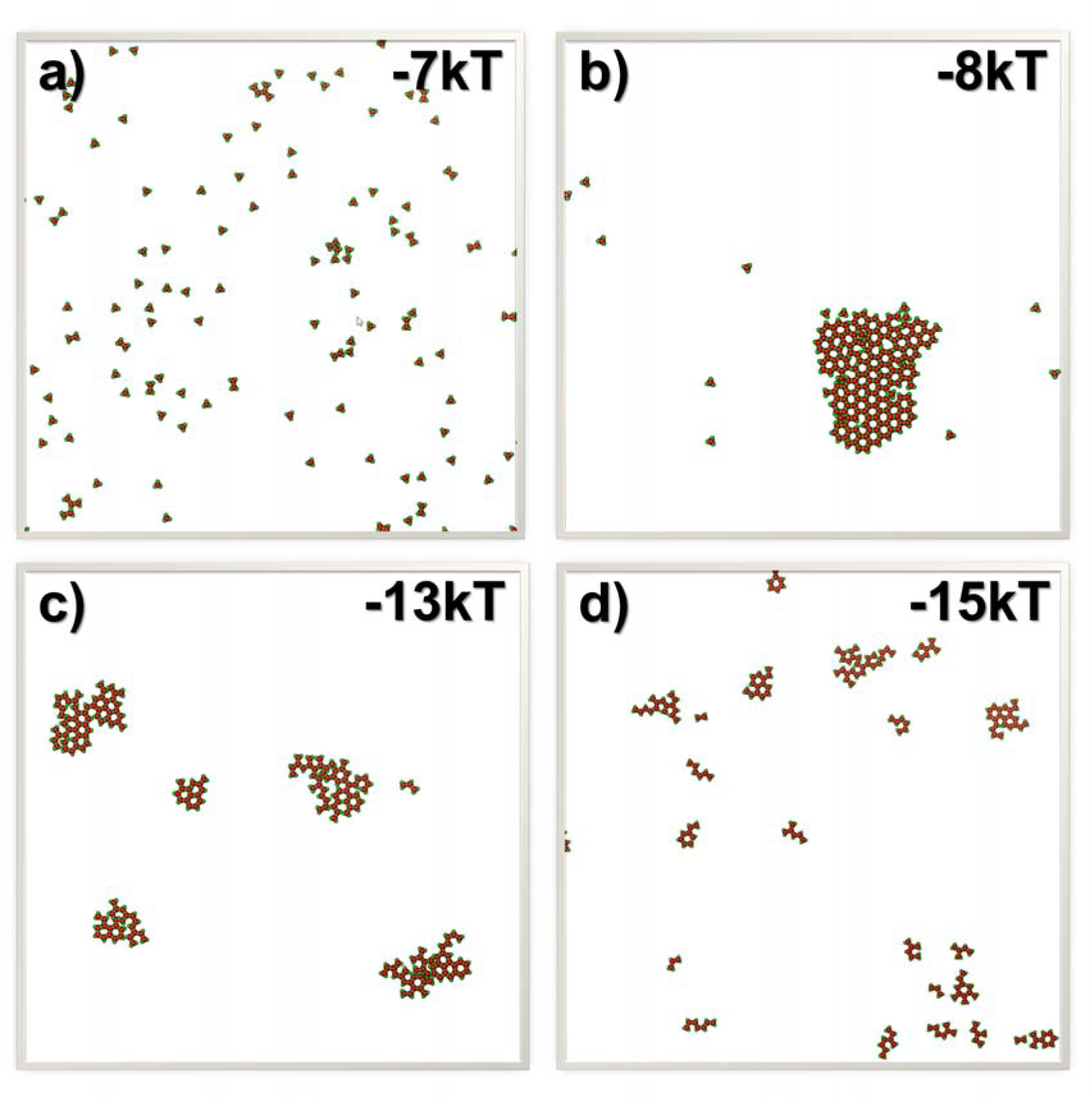
To illustrate the relation between the spatial properties of TRAF clusters and the strength of their *cis*-interactions, the final configurations from the systems which binding affinities equal -7kT and -8kT are plotted in **(a)** and **(b)**. These two plots show that the clustering can generate very different spatial pattern with only a slight change in the binding affinity. On the other hand, the final configurations from the systems which binding affinities equal -13kT and -15kT are plotted in **(c)** and **(d)**, respectively. These two plots suggest that systems with stronger binding affinity contain more clusters with smaller sizes.

Taken together, our simulation results show that the increase of *cis-*binding affinity triggers the phase transition of TRAF clustering. This phase transition results in a threshold-like signal response from the downstream pathway. Moreover, the large-scale TRAF clusters is kinetically slow to assemble, but highly stable after its formation. This slow kinetics of the assembling process and high stability of the clustered structure assure that the signaling pathway only responses to a persistent and high dose of extracellular stimulations, therefore promote the fidelity of signal transduction within stochastic cellular environments [46]. Finally, the oscillation of NF--κB signals can only be archived through a very dynamic process by *cis*-interactions with moderate binding affinities. The TRAF proteins with overly strong *cis*-interactions will be kinetically trapped in small clusters, and further impede their function in activating the downstream signal oscillation. This suggests that the molecular interactions in the NF-κB signaling pathway are tuned within a specific range by natural selection to maintain its appropriate functions.

## Conclusions

NF-κB signaling pathway is one of the most important cell signaling pathways involved in inflammatory responses [47]. Recent experimental evidences started to show that signal transduction in the pathway is spatially modulated by its different molecular components [48]. However, the molecular mechanism of these spatial regulation and their functional impacts are not well understood. In order to tackle this problem, we developed a hybrid model in which the NF-κB signaling pathway is decomposed into two simulation scenarios. The physical process of TRAF clustering at membrane proximal region is simulated by a rigid-body-based diffusion-reaction algorithm, while the downstream signaling network is simulated by stochastic simulation with Gillespie algorithm. These two algorithms are further synchronized under a multiscale simulation framework. Using this simulation method, we illustrated that the formation of TRAF-mediated 2D signaling platform is a critical factor to regulate the downstream oscillatory dynamics in the signaling network. The modification of *cis*-interaction between TRAF proteins leads to the changes of their clustering patterns at membrane proximal regions, and further affects the NF-κB response. Interestingly, our results show that mutations either weaken or strengthen this *cis*-interaction can cause the abolishment of oscillation in the pathway. This observation suggests that molecular elements and their interactions in a signaling network are elaborately designed to carried out their appropriate functions. In summary, the hybrid simulation developed in this study shows possibility to model a signaling system with both spatial resolution and functional implication. The results from the simulations provides the general biological insights to the interplay between the spatial assembly of individual signaling molecules and the threshold-like output from the entire signaling network, which offers a potentially new way to control signaling pathways in cellular systems.

## Methods

### Mathematical representation of the signaling network

Following the spatial assembly of TRAF signaling platform, the changes of population for each type of molecular components in the downstream signaling network are quantitatively described by a set of ordinary differential equations (ODE). Specifically, the numbers of active IKK and inactive IKK in the system at time *t* are changed by solving the following two equations.

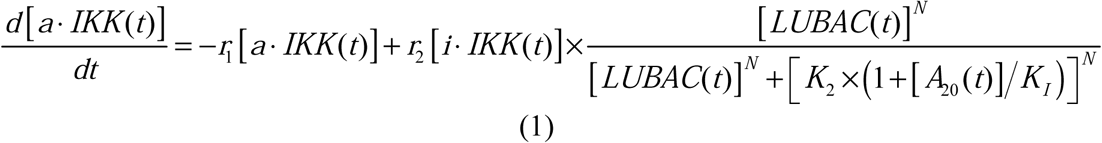

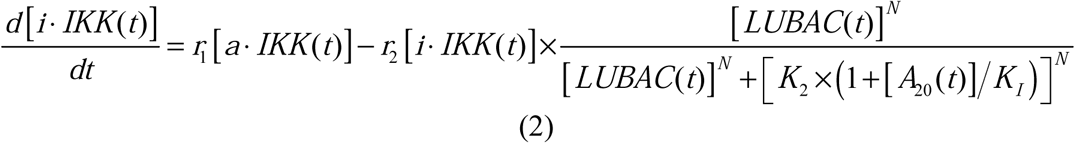

In above equations, *[a·IKK(t)]* and *[i·IKK(t)]* are the numbers of active IKK and inactive IKK at time *t*, respectively. The first term on the right-hand side of equation (1) represents the transition of IKK from its active form to inactive form, where the parameter *r*_*1*_ indicates the rate of the transition. The second term on the right-hand side of equation (1) represents the transition of IKK from its inactive form to active form. This reaction is catalyzed by the upstream poly-ubiquitin chains LUBAC, in which *r*_*2*_ and *K*_*2*_ are the maximal rate and saturation coefficient in the catalysis. The number of poly-ubiquitin chains in the system *[LUBAC(t)]* is determined by the number of *cis*-interactions between TRAF trimers in the upstream signaling platform. Moreover, considering that LUBAC itself is a highly ordered signaling machinery, a Hill coefficient [49] *N* is used to model the cooperativity in its assembly and its recruitment of IKK. Finally, the catalysis of this transition reaction is also restrained by protein A_20_ as a competitive inhibitor. The efficiency of inhibition is controlled by the concentration of A_20_ and the parameter *K*_*I*_.

The activated IKK kinase further phosphorylates the inhibitory IκB subunit in the IκB/*NF-*κ*B* complex. The number of phosphorylated complexes in the system at time *t* can be changed by solving the following equation.

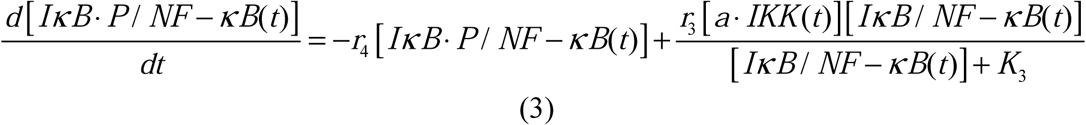

In above equation, *[I*κ*B/NF-*κ*B(t)]* and *[I*_κ_*B·P/NF-*_κ_*B(t)]* are the numbers of phosphorylated and un-phosphorylated complex at time *t*, respectively. The second term on the right-hand side of equation (3) describes the phosphorylation process, which rate depends on the concentration of activated IKK *[a·IKK(t)]*. The parameters *r*_*3*_ and *K*_*3*_ in the reaction are the maximal rate and saturation coefficient in the phosphorylation as described by Michaelis-Menten kinetics. The first term on the right-hand side of equation (3) describes the dissociation reaction of IκB/NF-κB complex due to the phosphorylation of IκB, in which *r*_*4*_ represents the rate constant of dissociation.

On the other hand, the number of unphosphorylated complex in the system is decreased by the IKK induced phosphorylation, but increased by the association reaction between monomeric IκB and NF-κB, as described by the following equation.

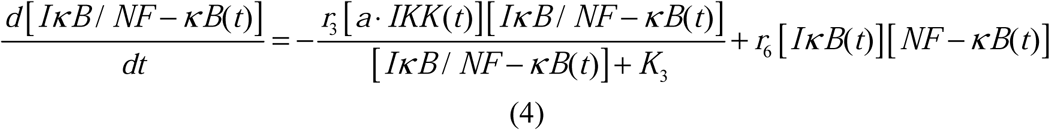

The parameter *r*_*6*_ in equation (4) is defined as the rate constant which regulates the association between monomeric IκB and NF-κB in cytoplasm. Their numbers at time *t, [I*_κ_*B(t)]* and *[NF-*_κ_*B(t)]* are changed by the following two equations:

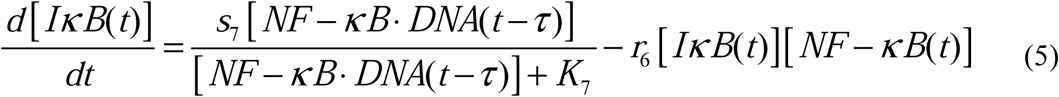

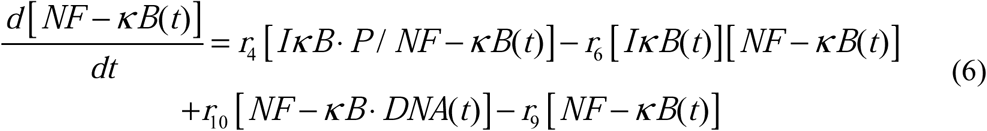

The second term on the right-hand side of equation (5) delineates the dissociation of IκB/NF-κB complex, as described above. Additionally, the dissociation of NF-κB from phosphorylated IκB enters cell nucleus and binds to its targeted DNA sequence as a transcription factor. The formation of NF-κB·DNA complex regulates the expression of specific genes, including its own inhibitor IκB, which corresponds to the first term on the right-hand side equation (5). We introduced the constant _τ_ in the reaction to describe the time delay between NF-κB·DNA formation and IκB expression. We assume that the expression rate of IκB depends on the concentration of activated transcriptional factors *[NF-*_κ_*B·DNA(t-*_τ_)] and reaches a maximum value *s*_*7*_. The parameter *K*_*7*_ is the saturation coefficient in the regulation of protein synthesis as described by Michaelis-Menten kinetics. In equation (6), on the other hand, the first two terms on its right-hand side have been introduced above, while the next two terms described the dynamic process in which NF-κB is translocated from cytoplasm into cell nucleus and binds to its target gene. Correspondingly, the parameters *r*_*9*_ and *r*_*10*_ indicate the rates for the association and dissociation of the NF-κB·DNA complex, respectively. Given these two rate constants, the number of the NF-κB·DNA complex in the system at time *t* can be changed by solving the following equation.

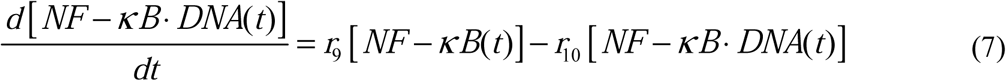

While the NF-κB dissociated from the IκB·P/NF-κB complex can either bind to DNA or reassociate with unphosphorylated IκB, the phosphorylated IκB that is dissociated from the IκB·P/NF-κB complex will be degraded with a rate constant d5, as described by the following equation.

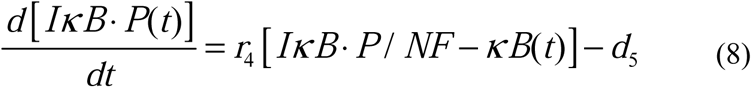

Finally, the change of protein A_20_ is defined by the following equation.

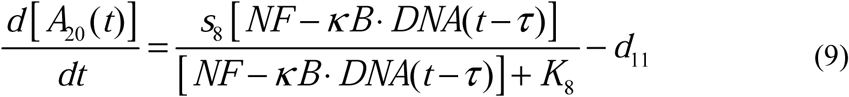

Similarly, its expression is regulated by the NF-κB·DNA complex, as represented the first term on the right-hand side equation (9). The same constant _τ_ is introduced to describe the time delay between NF-κB·DNA formation and A_20_ expression. The parameters *s*_*8*_ and *K*_*8*_ in the catalysis of protein synthesis give the maximal rate and saturation coefficient of the reaction as described by Michaelis-Menten kinetics. The second term on the right-hand side of the equation determines the degradation rate of A_20_.

### Numerical algorithm of the hybrid simulation

The dynamics of the signaling pathway is simulated by two coupled modules. The oligomerization of TRAF scaffold proteins is simulated by a diffusion-reaction algorithm, while the downstream signaling events are simulated by the Gillespie algorithm. In the diffusion-reaction algorithm, we assume the TRAF trimers have bound to the upstream ligand-receptor complexes. Therefore, their movements are confined within the membrane proximal area, which is modeled by a layer of two-dimensional flat surface. Each TRAF protein, together with its upper-bound ligand-receptor complex, is modeled as a single rigid body. In order to capture the basic structural information of the trimeric scaffold protein, the rigid body further contains four connecting groups. The three TRAF-C domains and coiled-coil regions are placed at the center, surrounded by three other groups representing their RING domains. The angle between every two surrounding groups is 120°. Each surrounding group also contains a binding site to mimic the *cis*-interaction between RING domains.

Given the model representation, the diffusion-reaction simulation is started from an initial configuration, in which a large number of TRAF rigid bodies are randomly distributed within the two-dimensional membrane-proximal region. Each time step in the simulation after the initial configuration is broken down into two scenarios [50]. During the first scenario, all TRAF trimers are chosen by random order to undergo diffusions within the two-dimensional area. The rotations of each trimer are restricted along the surface normal, while the translational movements are along any direction in the plain. The periodic boundary condition is applied so that any trimer leaving the 2D simulation box will enter its opposite side. Following the scenario of molecular diffusions, the binding kinetics of cis-interaction between TRAF trimers is simulation in the second scenario. The association between two TRAF trimers can be triggered if: 1) the distance between any binding sites within these two trimers is below a predetermined cutoff; and 2) their packing angle equals 180°, as shown in **Figure 1a**. If a *cis*-interaction is formed between two trimers, they will stop diffusing to facilitate further oligomerization. On the other hand, dissociation of a *cis*-interaction occurs with a probability that is determined by its binding affinity. After dissociation, two trimers can either reassociate as a geminate recombination if the distance and packing angle between any pair of their binding sites satisfy the association criteria, or diffuse farther away from each other. The new configuration is updated at the end of each simulation time step after both diffusion and reaction scenarios are sequentially performed. Finally, the iteration of above diffusion-reaction process will not be terminated until the dynamics of the simulated system reaches equilibrium.

The high-order clusters formed by TRAF scaffold proteins provide a platform to recruit the poly-ubiquitin chains. The downstream signaling processes activated by the poly-ubiquitination are modeled by non-spatial stochastic simulation algorithm. This algorithm was developed by Gillespie in order to study biochemical reactions [51]. Given the initial condition of a signaling network, the populations of each molecular species in the system are propagated in a digitalized and stochastic manner. In detail, within each simulation step, the rates for all reactions are sequentially calculated by the mathematical formulas described in the previous section of the method. A cumulative distribution function is generated by adding up all these rates. A random number is used to select one of these reactions based on the cumulative distribution function. Populations for all species are then updated according to the selected reaction. The simulation moves forward iteratively by above procedure. This Gillespie algorithm has further been coupled with the diffusion-reaction model [52], so that the spatial information of TRAF clustering can be integrated into the downstream signaling processes. Specifically, after the diffusion-reaction simulation, the number of newly formed *cis*-interactions between TRAF trimers can be obtained. Based on current knowledge about the function of TRAF protein in NF-κB signaling pathway, we assume that poly-ubiquitin chains are recruited at each *cis*-binding interface of TRAF trimers. Therefore, the number of LUBAC will be simultaneously updated. This new number of LUBAC enters the next step of the Gillespie simulation to guide the activation of IKK. In turn, the outputs from the Gillespie simulation affect the diffusion-reaction simulation as follow. Based on the Gillespie algorithm, the time of the simulation system is moved forward by a specific time interval τ. Given this new time interval, *n* steps of the diffusion-reaction simulation are carried out to generate a new spatial configuration. The number *n* is calculated as τ/Δt, in which Δt is the length of the diffusion-reaction simulation time step. This calculation is to ensure that the time scale between spatial processes of TRAF clustering and the downstream signaling pathway can be synchronized.

We described a generic framework of NF-κB signaling network in which some factors have not been simulated before. It is not realistic to derive consistent model parameters from various experimental measurements or previous computational models. As a result, the values of all parameters were chosen on a heuristic basis from the biologically meaningful range, so that the oscillatory nature of the pathway under a persistent ligand stimulation can be qualitatively reproduced within the timescale that is close to the experimental observation. The values of these parameters in the simulation are listed in the supplemental **Table S1**. The variations in these parameters will not significantly affect the general dynamic patterns of the system. It is more important to recognize how the spatial-temporal dynamics of the signaling system is quantitatively modified by a small perturbation in the parameters space.

## Acknowledgement

This work was supported by the National Institutes of Health under Grant Numbers R01GM120238 and R01GM122804. The work is also partially supported by a start-up grant from Albert Einstein College of Medicine. Computational support was provided by Albert Einstein College of Medicine High Performance Computing Center.

## Author Contributions

K.D. and Y.W. designed research; K.D. performed research; K.D. and Z.S. analyzed data; K.D. and Y.W. wrote the paper.

## Data Availability

Data sharing is not applicable to this article as no datasets were generated or analyzed during the current study.

---

**Table 1:** the information about the simulation parameters in the signaling network

| Index | Description | Mathematical Representation | Parameter | Value | Remark |
| --- | --- | --- | --- | --- | --- |
| 1 | Deactivation of IKK | $r_1 [a \cdot IKK(t)]$ | $r_1$ | $0.2s^{-1}$ | Rate of transition |
| 2 | Activation of IKK | $r_2 [i \cdot IKK(t)] \times \frac{[LUBAC(t)]^N}{[LUBAC(t)]^N + [K_2 \times (1 + [A_{20}(t)]/K_I)]^N}$ | $r_2$ | $0.4s^{-1}$ | Maximal rate of catalysis |
| | | | $K_2$ | 240nM | Saturation coefficient of catalysis |
| | | | $K_I$ | 10nM | Inhibition coefficient of catalysis |
| | | | $N$ | 25 | Hill coefficient |
| 3 | Phosphorylation of IκB | $\frac{r_3 [a \cdot IKK(t)] [I\kappa B / NF - \kappa B(t)]}{[I\kappa B / NF - \kappa B(t)] + K_3}$ | $r_3$ | $0.08s^{-1}$ | Maximal rate of catalysis |
| | | | $K_3$ | 500nM | Saturation coefficient of catalysis |
| 4 | Dissociation of IκB/NF-κB complex | $r_4 [I\kappa B \cdot P / NF - \kappa B(t)]$ | $r_4$ | $0.4s^{-1}$ | Dissociation rate |
| 5 | Degradation of IκB | $d_5$ | $d_5$ | $0.8nMs^{-1}$ | Degradation rate |
| 6 | Association of IκB/NF-κB complex | $r_6 [I\kappa B(t)] [NF - \kappa B(t)]$ | $r_6$ | $0.008nM^{-1}s^{-1}$ | Association rate |
| 7 | Synthesis of IκB | $\frac{s_7 [NF - \kappa B \cdot DNA(t - \tau)]}{[NF - \kappa B \cdot DNA(t - \tau)] + K_7}$ | $s_7$ | $0.8s^{-1}$ | Maximal rate of synthesis |
| | | | $K_7$ | 50nM | Saturation coefficient of synthesis |
| 8 | Synthesis of A<sub>20</sub> | $\frac{s_8 [NF - \kappa B \cdot DNA(t - \tau)]}{[NF - \kappa B \cdot DNA(t - \tau)] + K_8}$ | $s_8$ | $0.4s^{-1}$ | Maximal rate of synthesis |
| | | | $K_8$ | 100nM | Saturation coefficient of synthesis |
| 9 | Association of NF-κB with DNA | $r_9 [NF - \kappa B(t)]$ | $r_9$ | $0.8s^{-1}$ | Transition rate |
| 10 | Dissociation of NF-κB from DNA | $r_{10} [NF - \kappa B \cdot DNA(t)]$ | $r_{10}$ | $0.08s^{-1}$ | Transition rate |
| 11 | Degradation of A<sub>20</sub> | $d_{11}$ | $d_{11}$ | $0.1nMs^{-1}$ | Degradation rate |

